# Sub-nucleolar trafficking of Hendra virus matrix protein is regulated by ubiquitination and oligomerisation

**DOI:** 10.1101/2023.08.10.552741

**Authors:** Stephen M. Rawlinson, Tianyue Zhao, Florian A. Gomez, Cassandra T. David, Christina L. Rootes, Patrick F. Veugelers, Ashley M. Rozario, Cameron R. Stewart, Toby D.M. Bell, Gregory W. Moseley

## Abstract

Hendra virus (HeV) is a highly pathogenic member of the Henipavirus genus (order *Mononegavirales*), the replication cycle of which occurs primarily in the cytoplasm. The HeV matrix protein (HeV M) plays critical roles in viral assembly and budding at the plasma membrane, but also undergoes nuclear/nucleolar trafficking, to accumulate in nucleoli early in infection and, later, localise predominantly at the plasma membrane. Previously we found that HeV M protein targets specific sub-nucleolar compartments (corresponding to the FC-DFC (fibrillar centre (FC)/dense fibrillar component (DFC)) where it interacts with the nucleolar protein Treacle and modulates rRNA biogenesis by subverting the host nucleolar DNA damage response, indicating the importance of specific sub-nucleolar trafficking to infection. However, the mechanisms underlying targeting and movement between sub-nucleolar compartments by viral or cellular proteins remain poorly defined. Here, we assessed the molecular regulation of HeV M protein nucleolar/sub-nucleolar trafficking, finding that in infected cells and in cells expressing HeV M protein alone, M protein localizes into Treacle-enriched FC-DFC at early time points, and that FC-DFC localization is subsequently lost due to relocalization into the surrounding granular component (GC) of the nucleolus. Analysis using mutated M proteins and pharmacological modulation of ubiquitination indicate that this dynamic localization is regulated by ubiquitination and oligomerisation, with ubiquitination required for retention of HeV M in Treacle-enriched sub-nucleolar compartments, and oligomerisation required for egress. To our knowledge, this study provides the first direct insights into the dynamics and mechanisms of viral protein trafficking between sub-nucleolar compartments, important to the interplay between HeV M protein and host cell factors during infection.

**AUTHOR SUMMARY:** Henipaviruses, including Hendra (HeV) and Nipah viruses, cause deadly diseases in humans and livestock and are considered priority diseases by the World Health Organization due to their epidemic potential and lack of effective treatments. Understanding how these viruses interact with host cells is essential for developing new therapeutics. Our study examines the matrix (M) protein of henipaviruses and its interaction with the nucleolus, a cell structure that mediates ribosome production, and is a common target for various viruses, although their functions are largely unresolved. Previously, we showed that the HeV M protein targets a sub-nucleolar structure, called the FC-DFC, to modulate ribosome biogenesis. Here, we report that the M protein’s movement between sub-nucleolar compartments is controlled by two processes: ubiquitination, which causes accumulation of the protein in the FC-DFC, and oligomerization, which is associated with exit. Similar mechanisms are also observed in other henipaviruses. Our findings reveal mechanisms regulating the hijacking of host cell functions by henipaviruses and suggest new potential targets for antiviral therapies. This study is the first to investigate how viral proteins move within the nucleolus, offering new insights into interactions that may be significant to multiple viruses.

## INTRODUCTION

The nucleolus comprises a highly multifunctional structure with long established roles in ribosome biogenesis, as well as roles in cell cycle regulation, the DNA damage response (DDR), cellular stress responses, and signal recognition particle assembly [1, 2]. The nucleolus was recently shown to be a membrane-less organelle (MLO) comprising at least three immiscible liquid condensates that are formed by liquid-liquid phase separation (LLPS) [3]. The three components are the fibrillar centre (FC), dense fibrillar component (DFC) and granular component (GC). The FC is surrounded by the DFC to form functional units (FC-DFC), which are embedded within the GC (Fig. 1A) [4]. These compartments play distinct roles, including assembling a pipeline for the key steps of ribosome biogenesis.

**Figure 1.**
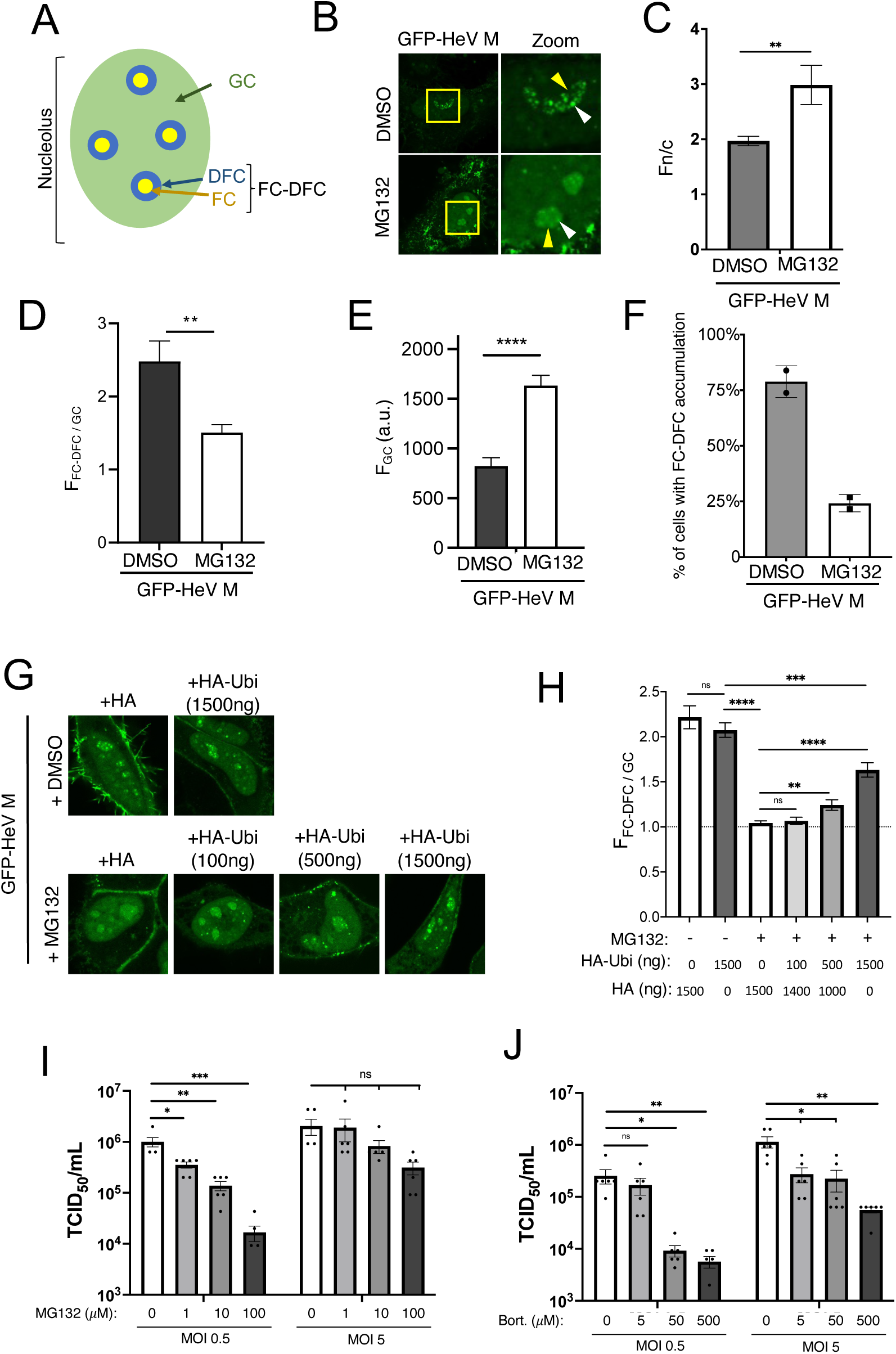
Ubiquitination regulates FC-DFC accumulation of HeV M and impacts on virus production. (A) Schematic of a nucleolus showing the three primary sub-compartments: fibrillar centre (FC), dense fibrillar component (DFC) and granular component (GC). The region composed of the FC and DFC compartments is referred to as the FC-DFC. (B) HeLa cells transfected to express GFP-HeV M protein were treated 18 h post transfection (p.t.) with MG132 or without (DMSO) for 6 h before CLSM analysis. Representative images are shown for each condition; yellow boxes are magnified in the zoom panel. Yellow arrowheads indicate nucleoli; white arrowheads indicate localization of M protein to sub-nucleolar compartments consistent with FC-DFC. Images such as those in B were analysed to determine: (C) the nuclear to cytoplasmic (Fn/c) fluorescence ratio; (D) the ratio of fluorescence of the FC-DFC to that of the GC (F_FC-DFC/GC_); (E) the fluorescence intensity of the GC (F_GC_) (arbitrary units (a.u.)), and (F) the % of HeV M-expressing cells with apparent accumulation in FC-DFC (histogram shows the percentage of M protein-expressing cells containing at least one nucleolus with evident accumulation of M protein into one or more FC-DFC). Histograms for C, D and E show mean ± S.E.M., n ≥ 24 cells for each condition (data from a single assay, consistent with two independent experiments); histogram in F shows mean percentage ± SD from two independent assays; ≥ 73 cells for each condition. (G) HeLa cells co-transfected with plasmid to express GFP-HeV M and with differing amounts of HA or HA-ubiquitin (HA-Ubi) expression plasmid (1500 ng total HA/HA-Ubi plasmid transfected, comprising HA-Ubi and/or HA, as indicated) and treated without (DMSO) or with MG132. (H) Images such as those in G were used to calculate the F_FC-DFC/GC_. Data from a single assay (n = 24 cells per sample), representative of two independent assays. (I, J) HeLa cells infected with HeV at MOI 0.5 or MOI 5 and treated with increasing concentrations of proteosome inhibitors, (I) MG132 or (J) Bortezomib (Bort), prior to collection at 42 h p.i. and determination of HeV titres (TCID50/ml ± S.E.M., n = 6). Statistical analysis used Student’s t-test; * p < 0.05; ** p < 0.01;*** p < 0.001; **** p < 0.0001; ns, non-significant.

Consistent with its multifunctionality, the nucleolus is a common target of diverse viruses [5–7]. This targeting is proposed to enable viral exploitation of diverse processes to usurp host cell biology and/or facilitate virus replication [5–7]. Despite the prevalence of viral protein nucleolar targeting, functional outcomes generally remain poorly understood. The potential nucleolar functions of viral proteins are of particular interest with respect to RNA viruses that typically have limited coding capacity and replicate their genomes in the cytoplasm, but nevertheless target specific proteins to nucleoli. These include the highly pathogenic non-segmented negative sense RNA viruses (nsNSVs) Hendra (HeV) and Nipah (NiV) viruses (genus *Henipavirus*, family *Paramyxoviridae*), the matrix (M) protein of which localizes to the nucleus and nucleolus during infection [8–11].

Henipavirus M proteins form stable homodimers, and play critical roles in virus assembly in the cytoplasm and in budding at the plasma membrane, which involves the formation of M protein oligomers [12, 13]. The subcellular localization of M protein is dynamic, being nucleolar early in infection before exiting the nucleolus/nucleus and accumulating at the plasma membrane for assembly/budding [8–11]. Interestingly, transit through the nucleolus is reported to be a prerequisite for M protein to fulfill assembly and budding functions, suggestive of a regulatory role of nucleoli in viral release [10, 11]. Genetic screens have indicated the importance of nucleolar proteins in infection, and proteomic datasets suggest that M protein interacts with multiple nucleolar proteins [8, 10, 14, 15]. However, potential intranucleolar roles of HeV M protein remained unresolved until the identification of a novel nucleolar function whereby HeV M localises to a sub-nucleolar compartment corresponding to the FC-DFC, where it interacts with Treacle protein and impairs ribosomal RNA (rRNA) biogenesis [14]. This process appears to be mediated by mimicry of a cellular process that normally occurs during a DDR. Thus subcellular trafficking underpins key functions of HeV M. However, how this trafficking is regulated, particularly between sub-nucleolar compartments and other regions of the cell remains unresolved. Indeed, the mechanisms regulating trafficking of proteins in general between sub-nucleolar liquid condensates is poorly understood, with no prior studies to our knowledge, for any viral protein.

Previously we showed that substitution of residue K258 in HeV M for alanine (HeV M K258A) impairs FC-DFC localisation/Treacle-binding and DDR modulation/budding activity, without preventing localization to the GC, where HeV M K258A accumulates [14]. K258 forms part of a bipartite nuclear localization sequence (NLS; often referred to as ‘NLS2’; M contains at least two NLSs; NLS1 is located at residues 82-87 [16, 17]) and is reported to be required for M ubiquitination (Fig. 2A) [10, 11]. It has been proposed that the M protein enters the nucleus via the NLS and accumulates within nucleoli before exiting the nucleolus and nucleus (mediated by a nuclear export sequence (NES)). Exit of the nucleolus/nucleus is reported to be triggered by ubiquitination requiring K258 by an unidentified ubiquitin ligase [18]. However, this model was proposed prior to the description of the functionally important localization of HeV M to sub-nucleolar compartments. As a result, the coordination and regulation of various trafficking steps, including trafficking within the nucleolus, remain undefined.

**Figure 2.**
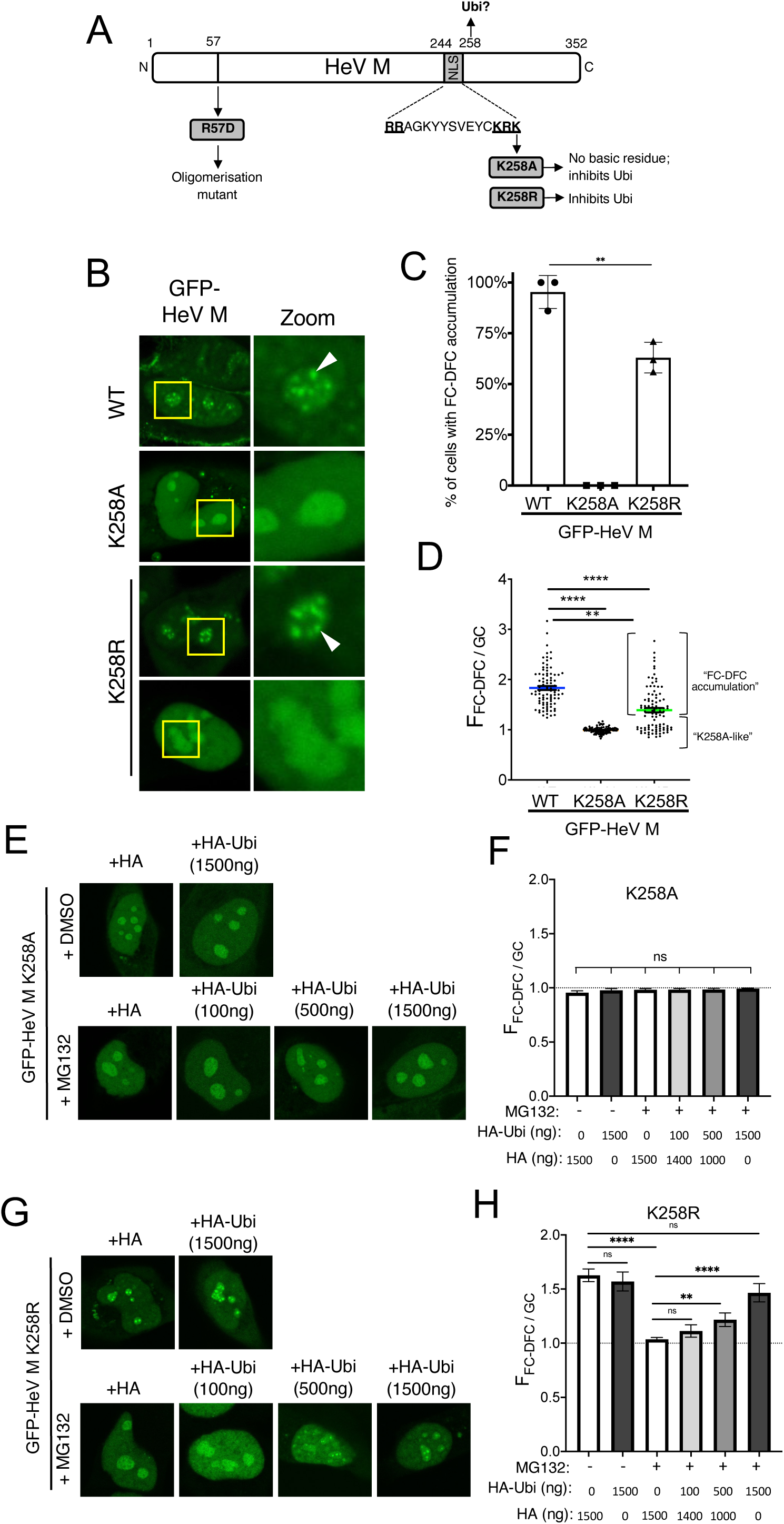
K258R mutation impacts sub-nucleolar trafficking of HeV M protein. (A) Schematic of the HeV M protein indicating the bipartite NLS (residues 244-258; critical basic residues are in bold and underlined) and residue K258, mutation of which regulates the ubiquitin status (Ubi) of M protein by removing the proposed ubiquitin site at 258, and affecting other sites. Mutations used in this study are indicated (grey boxes). (B) CLSM images of living HeLa cells expressing the indicated GFP-fused proteins (24 h p.t.); for HeV M WT and K258A, images are representative of 90-100% of cells in > 29 fields of view; for HeV M K258R two major populations (each representing c. 40-60% of the population) were observed, corresponding to either a “FC-DFC accumulation” (upper panel) or “K258A-like” (lower panel) phenotype. Nucleoli are highlighted by the yellow box, which is magnified in the zoom panel. White arrowheads indicate accumulation within FC-DFC. (C) Images such as those in B were analysed to determine the percentage of cells with clear FC-DFC accumulation of M protein (mean percentage ± SD, n = 3 separate assays, each sampling ≥ 119 cells). Student’s t-test with Welch’s correction was used to determine significance; ** p < 0.01.) (D) Images were analysed to determine F_FC-DFC/GC_ (mean ± S.E.M., n ≥ 90 cells for each condition, from three independent assays; green line in K258R indicates mean of all samples). The two distinct populations in the K258R sample are indicated. Comparison of K258R “WT-like” population with WT samples using Student’s t-test showed a significant difference (p < 0.01). (E-H) Images of HeLa cells co-transfected to express GFP-HeV M K258A (E) or K258R (G) with different amounts of HA or HA-ubiquitin-expressing plasmid imaged by CLSM (as in Fig. 1G, H). Images such as these were analyzed to determine the F_FC-DFC/GC_ of K258A (F) and K258R (H) expressing cells (n ≥ 17 cells for F, and n ≥ 21 for H; data from one assay, representative of two independent assays). Statistical analysis used Student’s t-test; * p < 0.05; ** p < 0.01; **** p < 0.0001; ns, non-significant.

The potential role of ubiquitination, and definition of other molecular mechanisms in sub-nucleolar localization are of particular interest as mechanisms regulating nucleolar/sub-nucleolar trafficking (which involves movement between LLPS structures) are poorly understood compared with those for nuclear trafficking, which involves conventional protein interactions with trafficking receptors and the nuclear pore complex. In this study, we examined the regulation of HeV M protein trafficking between the FC-DFC and GC finding that ubiquitination plays a crucial role. Interestingly, our data indicate that ubiquitination exerts opposing effects on sub-nucleolar and nucleocytoplasmic localization, suppressing exit from the FC-DFC to the GC while being required for egress from the nucleolus/nucleus. Furthermore, mutations affecting HeV M oligomerization indicated that oligomerization is necessary for FC-DFC egress. These findings provide new insights into the interplay of ubiquitination and oligomerization in HeV M protein nuclear/nucleolar transit and roles in modulation of host cell biology, viral assembly and budding.

## RESULTS

### Ubiquitination affects sub-nucleolar trafficking of HeV M protein

Previously HeV M protein was reported to be ubiquitinated at several sites, potentially including residue K258 (and equivalent residues in other henipaviruses), and that mutation of K258 to A or R (the latter preventing ubiquitination but retaining the positive charge) inhibits ubiquitination of several sites that were shown to be mono-ubiquitinated [10, 11]. Ubiquitination was implicated in M protein trafficking using the proteasome/ubiquitination inhibitor, MG132, which caused nuclear and nucleolar retention of HeV and NiV M proteins, and inhibited nuclear export and viral-like particle (VLP) production by NiV M proteins [10, 11].

To explore the possibility that ubiquitination regulates trafficking between sub-nucleolar condensates, we examined the effect of MG132 treatment on localization/accumulation of HeV M protein to sub-nucleolar punctate compartments (which correspond to FC-DFC; Fig. S1 and data previously reported data [14]) using confocal laser scanning microscopy (CLSM) analysis of living HeLa cells expressing GFP-fused wild-type (WT) HeV M (GFP-HeV M). GFP-HeV M accumulated with FC-DFC (Fig. 1B). Notably, FLAG-tagged HeV M protein also localised within FC-DFC (Fig. S2A) and interacted with Treacle (Fig. S2B), but not with the Treacle-binding mutant (K258A), as expected [14], suggesting that the GFP-tag is not causing artefacts.

As expected [10, 11], MG132 treatment resulted in an apparent increase in nuclear accumulation of GFP-HeV M protein (Fig. 1B); quantitative image analysis confirmed a significant increase in the nuclear to cytoplasmic fluorescence ratio (Fn/c) (Fig. 1C). This is consistent with previously reported impairment of nuclear export [10, 11]. In contrast, accumulation of M protein in the FC-DFC appeared to be reduced (Fig. 1B), and this effect was confirmed by a reduced ratio of fluorescence intensity of the FC-DFC compared with the GC (F_FC-DFC/GC_) (Fig. 1D), an increase in GC fluorescence indicative of movement of M protein from the FC-DFC into the surrounding GC (Fig. 1E), and a decrease in the number of cells with FC-DFC accumulation of M protein (Fig. 1F).

To confirm that the effects of MG132 on HeV M sub-nucleolar trafficking are due to ubiquitination, we co-transfected cells with a plasmid expressing HA-ubiquitin (HA-Ubi) to replenish ubiquitin depletion by MG132. Expression of HA-Ubi reversed the effect of MG132 in reducing FC-DFC accumulation in a dose-dependent fashion (Fig. 1G, H). These findings indicate that ubiquitination promotes the accumulation of HeV M protein within the FC-DFC, while reduced ubiquitination leads to its egress from the FC-DFC and accumulation in the GC. The observed increase in GC fluorescence is consistent with previous reports of an apparent enhancement of nuclear and nucleolar localization of M protein following MG132 treatment, proposed to reflect decreased export from the nucleus and corresponding decrease in egress from the nucleolus[10]. Our data suggest that the accumulation of diffuse (GC) fluorescence in the nucleolus following MG132 treatment is not solely due to reduced nuclear/nucleolar egress, but also increased egress from the FC-DFC to the GC. Notably, the opposing effects of ubiquitination on FC-DFC and nuclear/nucleolar GC localization indicate different mechanisms affecting trafficking between the compartments, such that FC-DFC localization is not simply the result of altered protein concentration in the nucleus/nucleolar compartment, but is specifically and distinctly regulated by ubiquitination.

Previously, proteosome inhibitors were shown to reduce NiV titers during live virus infections, indicating that ubiquitination plays a critical role in NiV infection [11]. To test if similar mechanisms occur during HeV infection, HeLa cells were infected with HeV at MOIs of 0.5 or 5, followed by treatment with proteosome inhibitors MG132 (Fig. 1I) and Bortezomib (Fig. 1J). Both inhibitors reduced virus titers in a dose-dependent manner at both MOIs, with statistically significant effects at both MOIs for Bortezomib and at MOI 0.5 MG132; the reduction observed for MG132 at MOI 5 was dose-dependent but not significant. These findings confirm the importance of ubiquitination in HeV infection, similar to what is observed with NiV [11].

### Conservative substitution of K258 to R reduces FC-DFC targeting by HeV M

We previously showed that K258A mutation in HeV M protein (HeV M K258A) abolishes its targeting to the FC-DFC, resulting in accumulation within the GC and loss of binding to the FC-DFC-enriched protein, Treacle [14]. This suggested that K258 forms part of a targeting signal due to its positive charge and/or affects sub-nucleolar localization due to its ubiquitination [10, 11]. The above data (Fig. 1) indicate that ubiquitination is required for the retention of HeV M within the FC-DFC. Mutation at K258 is reported to affect mono-ubiquitination at several sites, which has been suggested to indicate ubiquitination likely occurs at K258 but also impacts on other mono-ubiquitination sites in M protein [11]. To examine whether the effects we observed on FC-DFC localization following MG132 relate to ubiquitination at K258 or associated sites, we compared the effects on subcellular localization of HeV M by the substitutions K258A (which removes the positive charge and the potential ubiquitination site) and K258R (which retains a positive charge but lacks the lysine of the potential ubiquitination site) (Fig. 2A). Previous studies on equivalent mutations in NiV M protein indicated that the positive charge is important to function of the nuclear localization sequence (NLS) and nucleolar accumulation, while ubiquitination regulates nuclear export [11]. However, no effects on sub-nucleolar localization was reported, although our data indicate that ubiquitination has opposing effects on nuclear/nucleolar accumulation and FC-DFC accumulation (above).

CLSM analysis of cells expressing GFP-fused HeV M WT, K258A or K258R variants (Fig. 2B), indicated sub-nucleolar accumulation of WT M protein in c. 90% of cells, consistent with localization to FC-DFC (Fig. 2C). As expected [14], HeV M K258A protein did not localize/accumulate within FC-DFC, but accumulated within the GC in 100% of cells (Fig. 2B). In contrast, HeV M K258R displayed an intermediate phenotype, with a substantial proportion of cells (c. 60%) showing FC-DFC accumulation similar to WT and the remainder lacking FC-DFC accumulation similar to K258A (Fig. 2B, C). Consistent with this, the F_FC-DFC/GC_ ratio for HeV M K258A (c. 1.0) was significantly lower than that for HeV M WT (c. 1.8), while HeV M K258R showed an intermediate phenotype (c. 1.4) (Fig. 2D). The reduced F_FC-DFC/GC_ for HeV M K258R resulted from the presence of a K258A-like sub-population (for which the F_FC-DFC/GC_ was equivalent to that for K258A) and the fact that the F_FC-DFC/GC_ for the population with apparent FC-DFC accumulation was significantly lower than the F_FC-DFC/GC_ for HeV WT protein (p < 0.01) (Fig. 2D); thus, even in cells where HeV M K258R localized to the FC-DFC, this localization was impaired compared with the HeV M WT protein. Overexpression of HA-Ubi, treatment with MG132, or a combination of these conditions did not result in any significant FC-DFC accumulation of HeV M K258A (Fig. 2E, F), consistent with the positive charge at residues 258 being essential for FC-DFC localization [14]. Interestingly, MG132-treatment of cells expressing GFP-HeV-M-K258R resulted in a significant reduction in FC-DFC accumulation to reach levels similar to HeV M K258A, and HeV M WT with MG132/HA (Fig. 1H). Expression of HA-Ubi reversed this effect (Fig. 2G, H). Together, these data imply that ubiquitination dependent on K258 is required for efficient FC-DFC localization, but that ubiquitination at other K258-independent sites (either in M protein or other cellular proteins), is also required.

### Dynamic localization of HeV M in sub-nucleolar compartments is regulated by K258

The effects of the K258 mutation on HeV M protein accumulation in FC-DFC suggest potential impacts on a targeting sequence and/or affinity for specific components within the FC-DFC. For NiV M protein, K258 is proposed to be part of a NLS, which typically consists of short stretches of basic residues. Several basic residues (R244, R245, R256, R257, K258 in NiV M) are highly conserved among henipavirus M proteins [10, 11]. Thus, K258 may play a crucial role in nuclear import through the NLS and contribute to an overlapping targeting sequence for nucleoli/sub-nucleolar FC-DFC [11, 17]. Previous data indicated that ubiquitination dynamically regulates the nuclear localization of HeV M [10]. Moreover, NiV M protein undergoes dynamic and temporal regulation of localization during infection, being nuclear/nucleolar early in infection before nucleolar exit/nuclear export, and eventual accumulation and budding at the plasma membrane [9, 11]. Similarly, we found that in HeV-infected cells, HeV M localizes to sub-nucleolar compartments (FC-DFC) early during infection (7 hours post-infection (p.i.)), but becomes more diffuse in the nucleolus (i.e. accumulated into the GC), and with greater nuclear accumulation at 24 hours p.i. (Fig. 3A), consistent with observations for GFP HeV M WT protein and observations of dynamic nuclear/nucleolar localization of M protein in NiV infected cells [11]. Thus, we speculated that the observed differences in FC-DFC localization between HeV M WT and the K258 mutants (assessed at 24 hours post-transfection in Fig. 1 and Fig. 2) might be attributable to the dynamic regulation of various M protein trafficking signals related to the changes in HeV M localization during infection.

**Figure 3.**
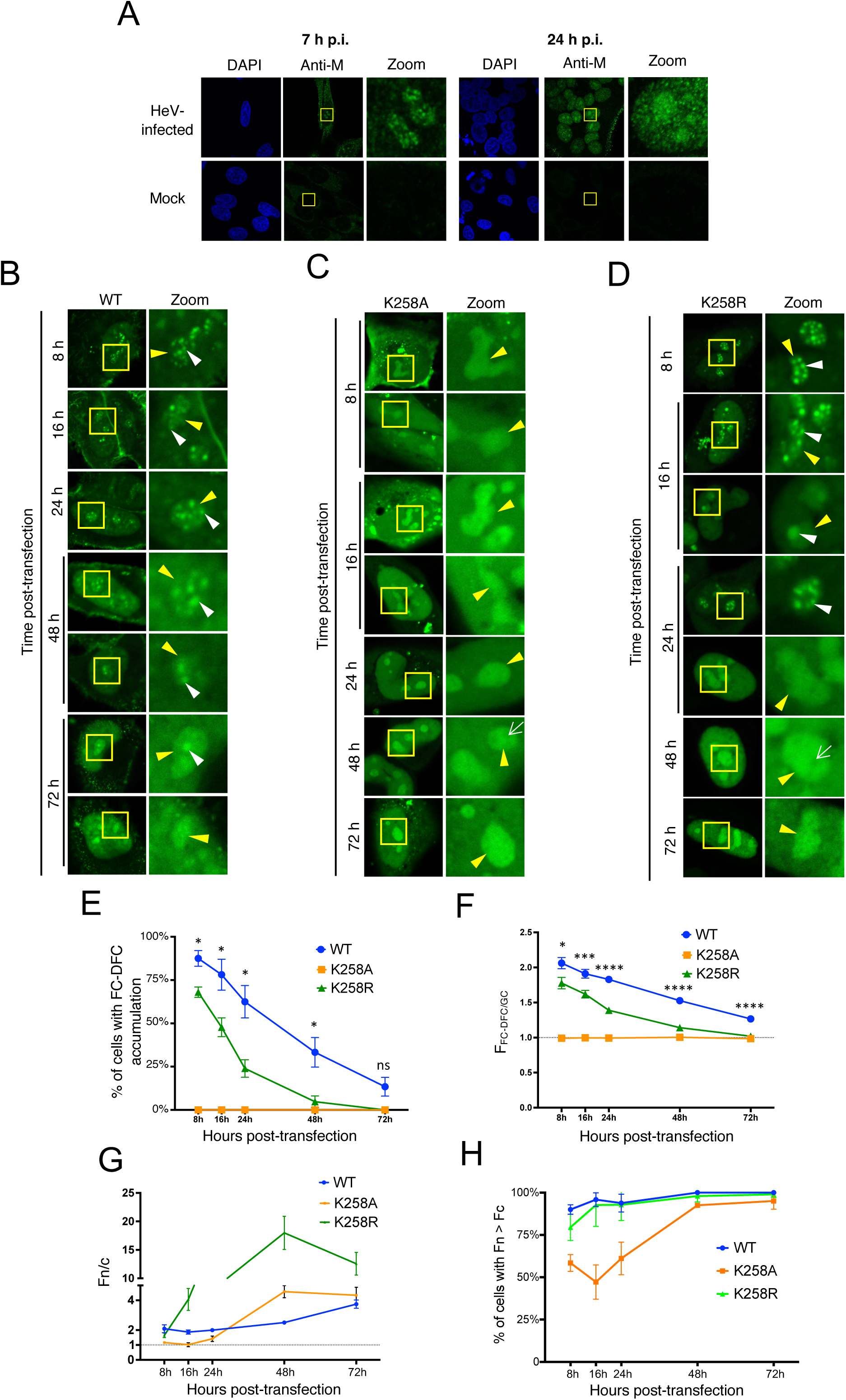
HeV M protein undergoes dynamic localization to the FC-DFC which is impacted by K258R mutation. (A) HeLa cells were mock-infected or infected with HeV (MOI 5) prior to fixation and immunostaining for HeV M protein at 7 h and 24 h post-infection (p.i.). Nuclei were detected using DAPI. Microscope settings and image correction are identical between equivalent mock and HeV infected images. (B-C) HeLa cells transfected to express the indicated proteins were analysed live at 8, 16, 24, 48 and 72 h p.t. by CLSM. Images representative of major phenotypes are shown for each condition; yellow boxes are magnified in the zoom panel. Yellow arrowheads indicate nucleoli; white arrowheads indicate accumulation of M protein in FC-DFC. Images such as those in B-C were analysed to determine: (E) the percentage of M protein-expressing cells with FC-DFC accumulation in any nucleolus in the cell (mean ± SD from three independent assays, n ≥ 59 cells for each condition); (F) F_FC-DFC/GC_ (mean ± S.E.M., n ≥ 55 cells for each condition from three independent assays, except for 8 h p.t. (WT, K258A, K258R), 16 h p.t. K258A, and 48 h p.t. WT samples, where data is from two assays; (G) Fn/c (mean ± S.E.M.; n ≥ 24 cells) and (H) percentage of cells with Fn > Fc (mean ± SD from three independent assays, except for 8 h p.t. (WT, K258A, K258R), 16 h p.t. K258A, and 48 h p.t. WT samples, where data is from two assays; n ≥ 23 cells for each condition). Student’s t-test was used to compare values for WT and K258R at each timepoint in D and E; * p< 0.05; *** p < 0.001; **** p < 0.0001; ns, non-significant.

To investigate this, we assessed the sub-nucleolar localization of HeV M WT and mutant proteins at time points from 8 h to 72 h p.t. (Fig. 3B-D). The WT HeV M protein exhibited clear accumulation in FC-DFC (typically multiple structures in each nucleolus) at 8 h p.t. (c. 85% of cells), which progressively diminished over the course of the experiment, accompanied by a more diffuse GC distribution with only around 10% of cells exhibiting accumulation of M protein in multiple FC-DFC at 72 h p.t. (Fig. 3E). This is consistent with a dynamic interaction whereby M protein initially enters the FC-DFC and then undergoes gradual egress to the GC. Measurement of the F_FC-DFC/GC_ confirmed a progressive loss of FC-DFC localization (Fig. 3F).

Consistent with roles of K258 in NLS activity of NiV M protein [10], the Fn/c for HeV M K258A was reduced compared with WT at 8 and 16 h p.t., supporting its involvement in nuclear import (Fig. 3G; Fn/c c. 2 for WT, compared with Fn/c c. 1 for K258A at both timepoints). Further analysis revealed that the reduction in nuclear localization of the K258A mutant was due to a significant proportion of cells with higher fluorescence intensity in the cytoplasm (Fc) than in the nucleus (Fn) at early time points, in contrast to cells expressing WT and K258R M protein (Fig. 3H; c. 50-60% of cells expressing K258A M protein showed Fn > Fc between 8-24 h p.t., whereas in cells expressing WT M protein, > 85% of cells showed Fn > Fc at all timepoints). However, over time, the K258A mutant gradually exhibited a proportion of cells with Fn > Fc similar to WT and K258R (nearly 100% of cells at 48 and 72 h p.t.), suggesting a delay in nuclear import of K258A compared to the other variants.

Despite reduced nuclear accumulation at early time points, HeV M K258A was strongly nucleolar at all time points, consistent with reduced nucleolar egress. However, no accumulation in FC-DFC was observed at any time point, and some images indicated absence of fluorescence from these structures (e.g. white arrow, 48 h p.t., Fig. 3C). Thus, HeV M protein can specifically partition between sub-nucleolar phase-separated compartments, dependent on K258, and this is independent of the accumulation in the nucleus, consistent with distinct mechanisms of trafficking/localization. Notably, similar dynamics were observed for WT NiV M, with the percentage of cells showing FC-DFC accumulation reducing over time (c. 25% of cells showing FC-DFC accumulation at 72 h p.t.), while NiV M-K258A remained excluded from FC-DFC at all timepoints (Fig. S3).

HeV M K258R accumulated to higher levels in the nucleus than the cytoplasm at early time points compared with K258A (Fig. 3D, upper panels and 3G), similar to WT M. This is consistent with a requirement for the positive charge in the NLS for efficient nuclear import, as reported for NiV M protein [11, 17]. Additionally, HeV M K258R accumulated to very high levels in the nucleus at later time points (48 h), consistent with an impaired nuclear export mechanism [10, 11]. HeV M K258R also showed clear FC-DFC localization in Treacle-enriched compartments at 8 h p.t. (Fig. 3D), similar to (but moderately reduced compared with) WT HeV M protein, followed by loss of FC-DFC localization over time. Thus, the presence of a basic residue at position 258 is necessary for initial entry and accumulation within the FC-DFC.

While FC-DFC localization of WT and K258R HeV M protein diminished following the initial accumulation, the apparent rate of loss was greater for HeV M K258R, such that by 24 h p.t. (Fig. 3E) c. 25% of HeV M K258R-expressing cells displayed FC-DFC localization compared with c. 60% for WT HeV M. By 48 and 72 h p.t. < 5% and 0%, respectively, of HeV M K258R-expressing cells displayed FC-DFC localization, and nucleoli with apparent exclusion from FC-DFC structures were apparent (e.g. 48 h p.t., Fig. 3D, white arrow, similar to observations for HeV M K258A. Calculation of the F_FC-DFC/GC_ ratio confirmed a significant decrease in FC-DFC localization by both HeV M WT and K258R over the course of the experiment, with a more rapid decrease for the latter (Fig. 3F). Thus, it appears that HeV M localises initially to the FC-DFC, dependent primarily on the presence of a positive charge at position 258. HeV M then relocalizes to the GC, and this process is accelerated in HeV M containing the K258R substitution that is impaired for ubiquitination, consistent with ubiquitination supporting retention into the FC-DFC.

### FC-DFC accumulation is enhanced by inhibition of HeV M protein oligomerization

Protein oligomerization has been associated with regulation of subcellular localization, including nuclear import and export of proteins such as p53 [19] and signal transducer and activator of transcription (STAT) proteins [20]. Moreover, oligomerization is implicated in the formation of MLOs through LLPS [21, 22]. Based on the crystal structure of HeV M protein, residue R57 of HeV M has been implicated in oligomerization *via* packing of HeV M dimers [13]. Mutation of R57 to D or E impairs the formation of VLPs by M protein, suggesting roles for oligomerization in the assembly and budding processes [13]. Assembly and budding is also dependent on K258 [10, 11], suggesting that nuclear and nucleolar localization/trafficking, and budding are intrinsically linked and/or involve coordinated regulation. Additionally, it has been hypothesized that ubiquitin modification at K258 may be necessary for M protein oligomerization [18]. However, roles of oligomerization in HeV M nuclear/nucleolar trafficking have not been investigated.

To examine this we analysed the localisation of HeV M WT or R57D as above, which indicated pronounced accumulation of HeV M R57D into the FC-DFC compared to the WT protein (Fig. 4A). The Fn/c was not significantly different between HeV M WT and R57D indicating similar function of the nuclear trafficking signals (Fig. 4B). However, the ratio of nucleolar to nuclear fluorescence (Fnu/n) (Fig. 4C) and F_FC-DFC/GC_ (Fig. 4D) was significantly increased for HeV M R57D compared to WT HeV M protein. Thus, R57D enhances the accumulation of HeV M in the FC-DFC, consistent with a role for oligomerization in regulating the localization of HeV M to nucleolar condensates. These data further indicate that the mechanisms underlying M protein FC-DFC localization differ from those regulating nuclear import/export. Interestingly, in contrast to results for WT M protein, MG132 treatment did not reduce FC-DFC accumulation (Fig. 4E, F). This suggests that the impaired ability of HeV M R57D protein to oligomerize both enhances FC-DFC localization, and causes it to become insensitive to the effects of reduced ubiquitination. These data are consistent with egress of M protein from the FC-DFC being dynamically regulated by ubiquitination, but also being dependent on the ability of M protein to undergo oligomerization.

**Figure 4.**
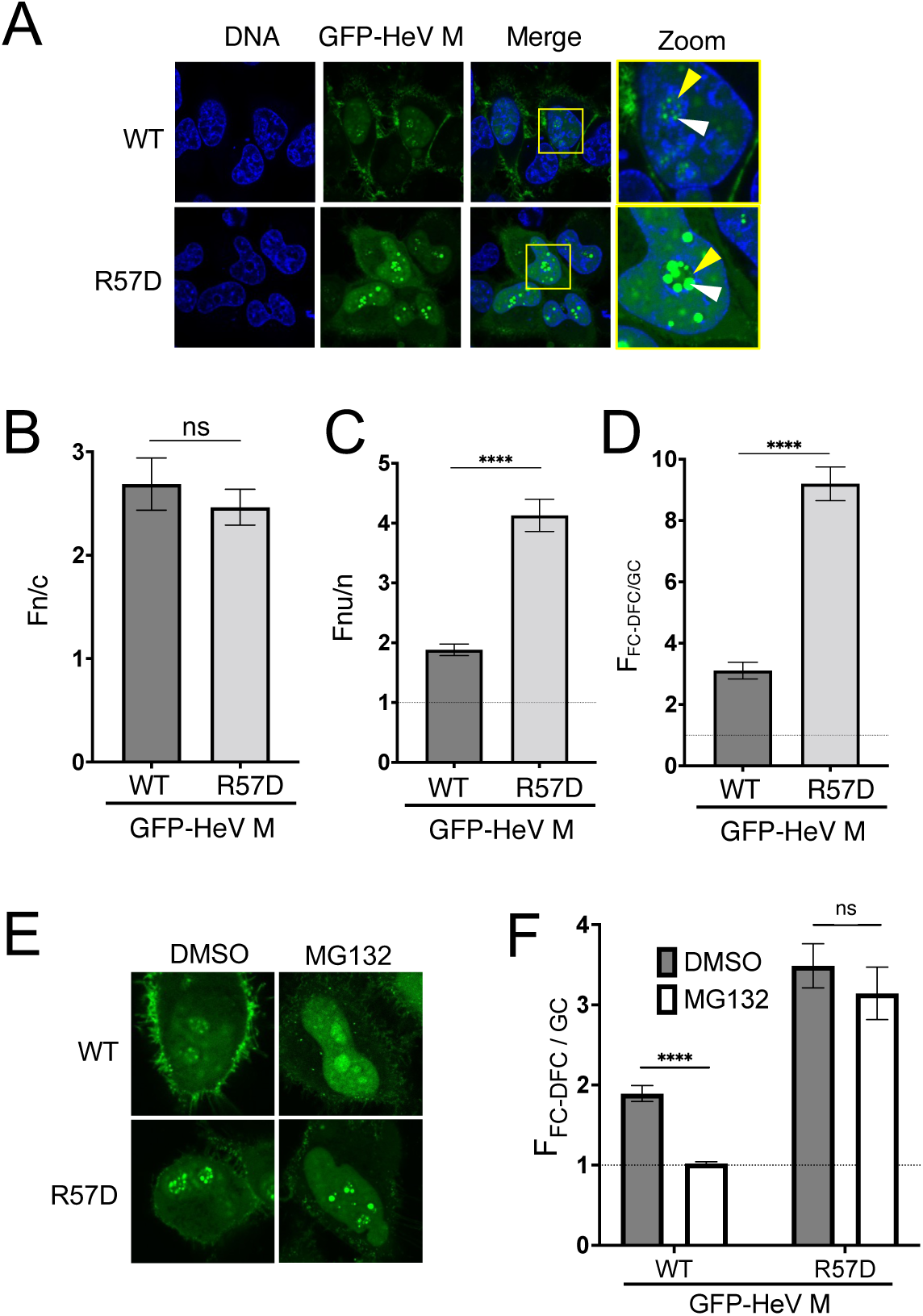
R57D mutation enhances accumulation of HeV M in FC-DFC and is insensitive to ubiquitination depletion. (A) CLSM images of live HeLa cells expressing the indicated proteins (24 h p.t.). Images are representative of 90-100% of cells in > 9 fields of view from a single assay, and typical of ≥ 4 independent assays. Hoechst 33342 (blue) was used to identify DNA/nuclei. Nucleoli are highlighted by the yellow box, which is magnified in the zoom panel. Yellow arrowheads indicate nucleoli; white arrowheads indicate accumulation in FC-DFC. (B-D). Images such as those shown in A were used to determine (B) Fn/c; (C) nucleolar to nuclear fluorescence ratio (Fnu/n; Fnu is the mean fluorescence intensity for the entire area of the nucleolus); (D) F_FC-DFC/GC_. B, C and D show mean ± S.E.M., n ≥ 24 cells for each condition. (E) CLSM images of HeLa cells expressing the indicated protein and treated without (DMSO) or with MG132. (F) Images such as those in E were analysed to determine F_FC-DFC/GC_ (n ≥ 14 cells from one assay; data representative of three independent assays). Statistical analysis used Student’s t-test; **** p < 0.0001; ns, non-significant.

### Accumulation of HeV M protein in the FC-DFC correlates with inhibitory function toward rRNA biogenesis and with binding to Treacle

HeV M protein accumulation in FC-DFC enables subversion of the nucleolar DDR pathway through targeting of Treacle. This function is strongly impaired by the K258A mutation [14]. R57D and K258R enhance (Fig. 4) and suppress (to a level intermediate between WT and K258A protein; Fig. 2), respectively, accumulation within the FC-DFC, suggesting that oligomerization/ubiquitination regulate functional FC-DFC interactions. To assess this directly, we used the Click-iT RNA Imaging Kit to measure the effects of M protein and mutants on rRNA biogenesis (Fig. 5A, B), and co-immunoprecipitation (co-IP) assays to assess interactions with Treacle (Fig. 5C), as previously described [14, 23].

**Figure 5.**
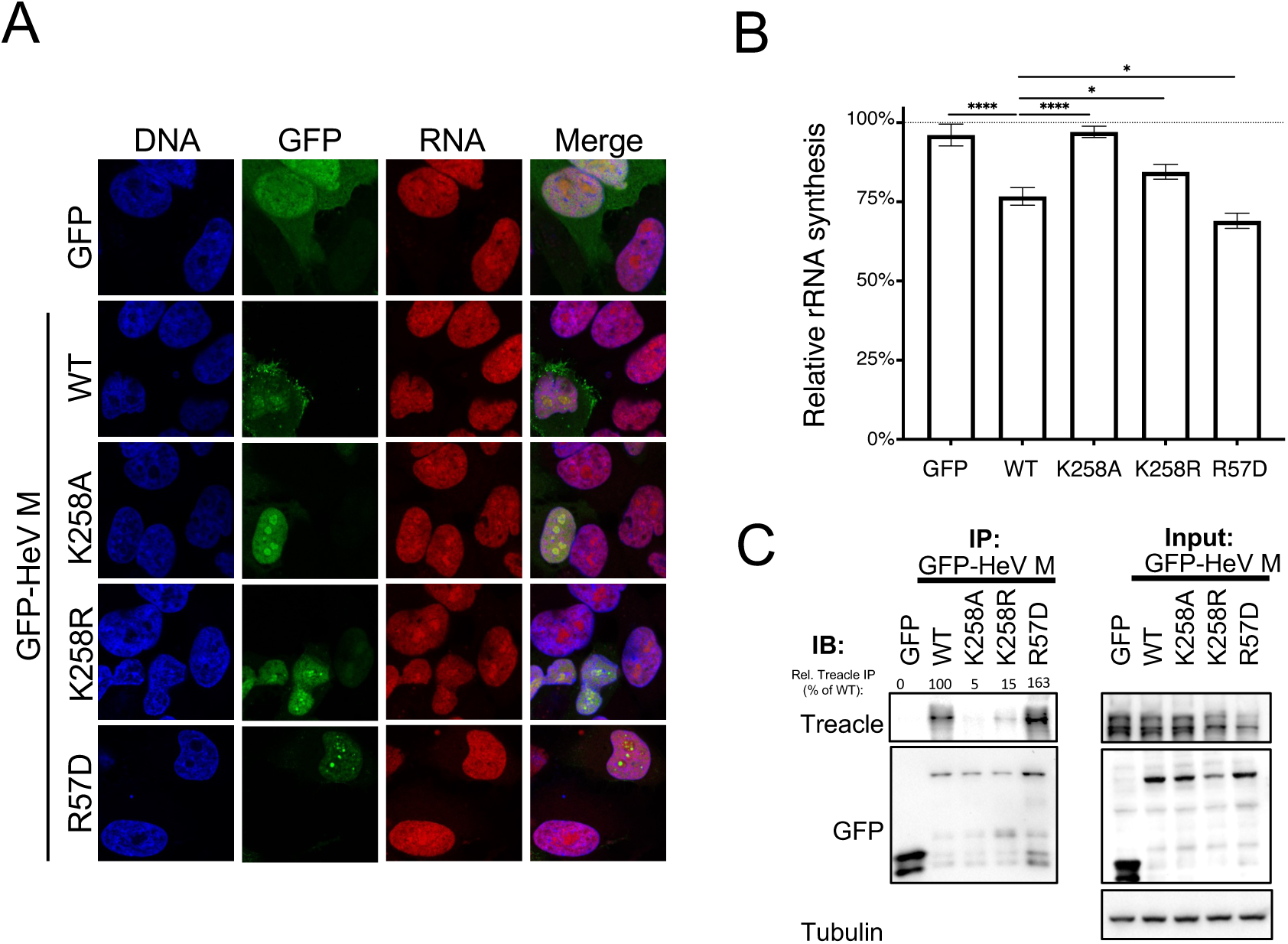
Silencing of rRNA synthesis and Treacle-binding by HeV M protein/mutants correlate with FC-DFC accumulation. (A) HeLa cells were transfected to express the indicated GFP-fused proteins before addition of EU reagent (23 h p.t.) for 1h, fixation (24 h p.t.), labeling to detect nascent RNA (EU fluorescence) and DNA (Hoechst 33342), and imaging by CLSM. (B) Images such as those in A were used to determine EU fluorescence in the nucleoli of GFP-positive cells, relative to that in non-GFP expressing cells in the same sample (mean relative EU fluorescence ± S.E.M., n ≥ 51 cells from two independent experiments). * p< 0.05; *** p < 0.001; **** p < 0.0001; ns, non-significant. (C) HEK-293T cells were transfected to express GFP or the indicated GFP-fused proteins prior to lysis (24 h p.t.) and IP for GFP. Cell lysate (input) and IP samples were subjected to IB using the indicated antibodies. Quantitation of Treacle band in IPs, relative to that detected in IP for GFP-HeV M WT are indicated.

Consistent with previous findings, WT HeV M protein, but not HeV M K258A significantly impaired rRNA biogenesis at 24 h p.t.; the level of inhibition was comparable with that observed for rRNA silencing by Treacle knockdown (c. 20-30% inhibition) [14, 24] (Fig. 5A, B). In contrast, R57D, significantly enhanced inhibition of rRNA biogenesis compared with WT M protein (c. 30% inhibition), consistent with strong FC-DFC localization and apparent loss of ubiquitin-regulated localization. The K258R mutation resulted in a phenotype for rRNA silencing that was intermediate between WT and K258A M proteins, correlating with their differing capacities to localize to the FC-DFC. Notably, the ability of WT M protein to inhibit rRNA biogenesis was lost by 72 h p.t. (Fig. S4), matching with dynamic loss of FC-DFC localization (Fig. 3). As expected, rRNA biogenesis in cells expressing HeV M K258A was unchanged between 24 and 72 h p.t., correlating with consistent lack of localization to the FC-DFC, while R57D was found to inhibit rRNA biogenesis at both 24 and 72 h p.t. (consistent with a sustained FC-DFC localization over time, see below and Fig. 6). Thus, the extent of FC-DFC accumulation appears to correspond with inhibition of rRNA biogenesis.

**Figure 6.**
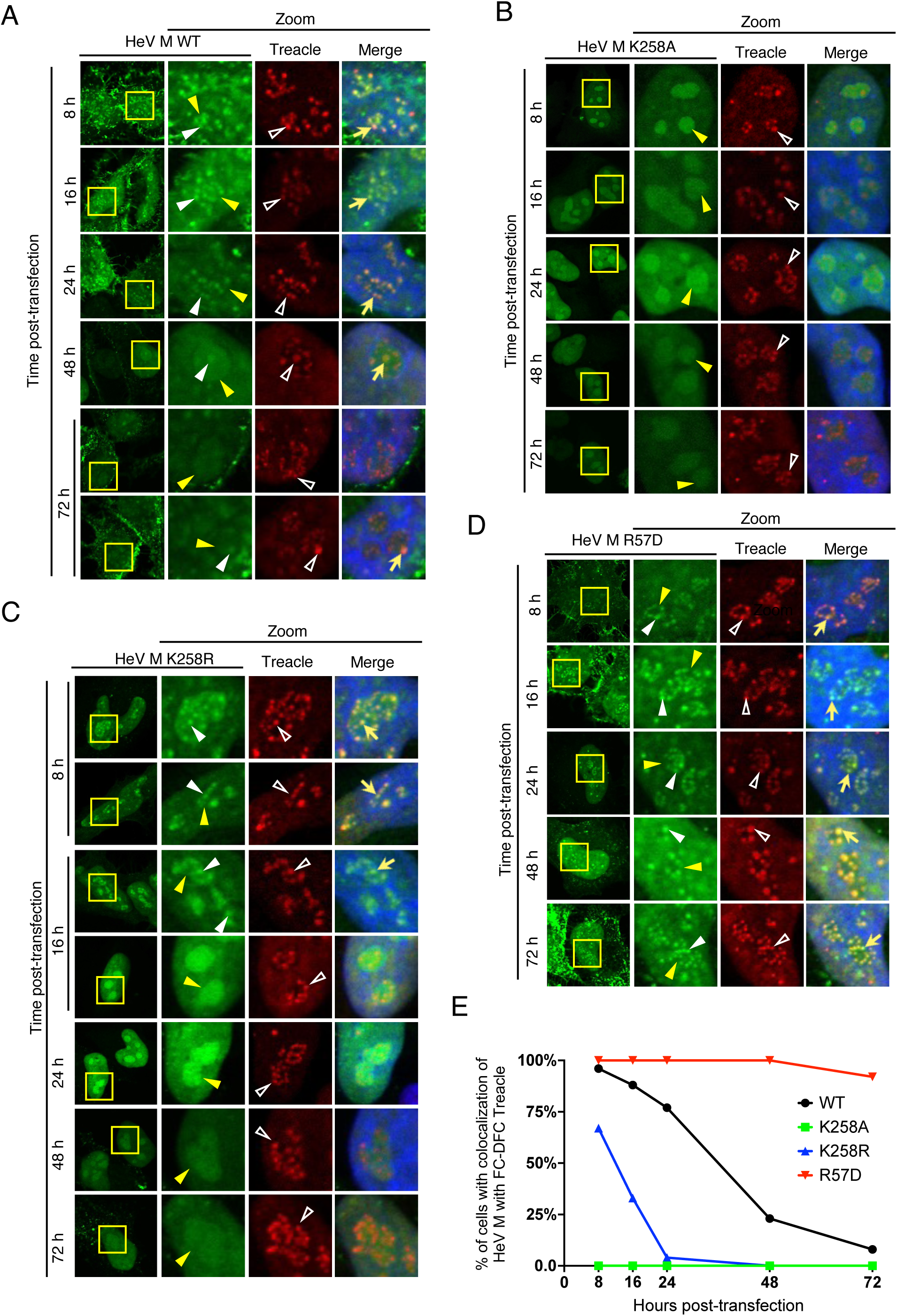
M protein FC-DFC accumulation decreases over time without loss of FC-DFC compartments. (A-D) HeLa cells were transfected to express the indicated proteins before fixation at 8, 16, 24, 48 and 72 h p.t. and immunostaining using anti-Treacle antibody (red) and imaging via CLSM. Hoechst 33342 (blue in Merge panels) was used to localize DNA/nuclei. Representative images are shown for each condition; yellow arrowheads indicate nucleoli; unfilled white arrowheads indicate Treacle in FC-DFC; white arrowheads indicate accumulation of M protein into FC-DFC; yellow arrows indicate colocalization of Treacle and HeV M protein in FC-DFC. (E) Images such as those in A-D were analysed to determine the percentage of cells expressing HeV M protein with evident colocalization of HeV M protein and Treacle in FC-DFC (n ≥ 23 cells for each condition).

Consistent with the differing localization and functional impact of the mutated proteins, co-IP and immunoblotting (IB) analysis indicated increased interaction of HeV M R57D protein with Treacle, compared with WT HeV M protein (Fig. 5C; c. 1.6-fold more Treacle precipitated with R57D). Furthermore, Treacle interaction of HeV M K258R was reduced compared with WT protein (c. 15% of WT levels), but was greater than HeV M K258A (c. 5 %), indicating an intermediate phenotype. Notably, Treacle is reported to form multiple isoforms [25], and only the upper band of Treacle appeared to IP with the M proteins (Fig. 5C), perhaps indicating interaction with specific Treacle isoforms. Taken together, these data indicate that FC-DFC accumulation, Treacle binding, and the inhibition of rRNA biogenesis by the HeV M protein are correlated, consistent with regulation of M protein function in rRNA production *via* ubiquitin and oligomerization dependent FC-DFC localization/Treacle interaction.

### Loss of HeV M FC-DFC accumulation does not relate to disruption or loss of FC-DFC

Loss of HeV M FC-DFC localization (Fig. 3) could be attributed to two possible mechanisms: (1) egress of the protein from intact FC-DFC structures, or (2) depletion of FC-DFC structures through events such as fusion or disassembly/disruption of the liquid bodies. To investigate these possibilities, we analysed cellular FC-DFCs directly by fixation and immunostaining of cells for Treacle at time points from 8-72 h p.t. to express HeV M WT, K258A, K258R, or R57D (Fig. 6A-D).

In cells expressing GFP-HeV M WT protein the appearance of FC-DFCs was similar throughout the experiment (Fig. 6A). At early time points, HeV M WT protein strongly colocalized with Treacle in FC-DFC, but this diminished over time (indicated by a reduced percentage of cells with detectable colocalization of HeV M and Treacle FC-DFCs), although multiple Treacle-enriched FC-DFCs lacking HeV M association remained detectable in nucleoli (Fig. 6A, E). Thus, HeV M protein appears to transit through FC-DFC, where it interacts with Treacle, before egress, with no significant disruption of FC-DFC structures. As expected, HeV M K258A showed no colocalization/accumulation in Treacle FC-DFC at any time point, despite the presence of multiple FC-DFC. HeV M K258R protein showed similar results to WT, but with more rapid egress, as expected (Fig. 3C), and no evident loss of FC-DFC (Fig. 6C, E). By 24-48 h p.t., co-localization was barely detectable, similar to K258A (Fig. 6E). In contrast, HeV M R57D exhibited clear colocalization with Treacle in all detectable FC-DFCs from 8 to 48 h p.t., with only a minor downward trend (colocalization with FC-DFC in c. 95% of cells measured) at 72 h p.t. (Fig. 6D, E). These findings are consistent with a requirement for oligomerization for HeV M protein to egress the FC-DFC, and with the capability of HeV M R57D to maintain efficient rRNA silencing function at 72 h p.t., by which time WT M protein has largely lost such function. Thus, it appears that HeV M protein transits through intact FC-DFC, with the loss of colocalization due to trafficking rather than disruption or major structural change to FC-DFC. These data further support that dynamic localization of M protein to FC-DFC, and interaction with Treacle, underlie specific silencing of rRNA biogenesis.

### HeV M proteins do not affect the structure or distribution of FC-DFC

The lack of major disruption of FC-DFC during HeV M protein accumulation or egress aligns with our previous findings [14]. Specifically, using single molecule localization microscopy (SMLM) with direct stochastic optical reconstruction microscopy (*d*STORM) of cells immunostained for Treacle, we found that HeV M WT protein accumulation in FC-DFC (24 h p.t.) does not cause ‘gross’ changes to FC-DFC numbers or to the structure of FC-DFC [14]. Based on the enhanced FC-DFC localization and function in rRNA silencing observed for HeV M R57D, we reasoned that, if M protein does have any structural effects on FC-DFC, they would be exaggerated by R57D. We thus analysed cells expressing GFP-HeV M WT, R57D or K258A protein, and cells expressing GFP alone or non-transfected cells, by immunostaining for Treacle followed by *d*STORM (Fig. 7A). Expression of HeV M WT, K258A (as previously [14]), K258R and R57D proteins had no significant impact on the size of Treacle-labelled FC-DFC (Fig. 7B). These data are consistent with specific intermolecular interactions of M protein with Treacle within FC-DFC regulating rRNA biogenesis, and the duration/extent of functional modulation being controlled by ubiquitination (and potentially multimerization)-dependent egress of M protein from FC-DFC.

**Figure 7.**
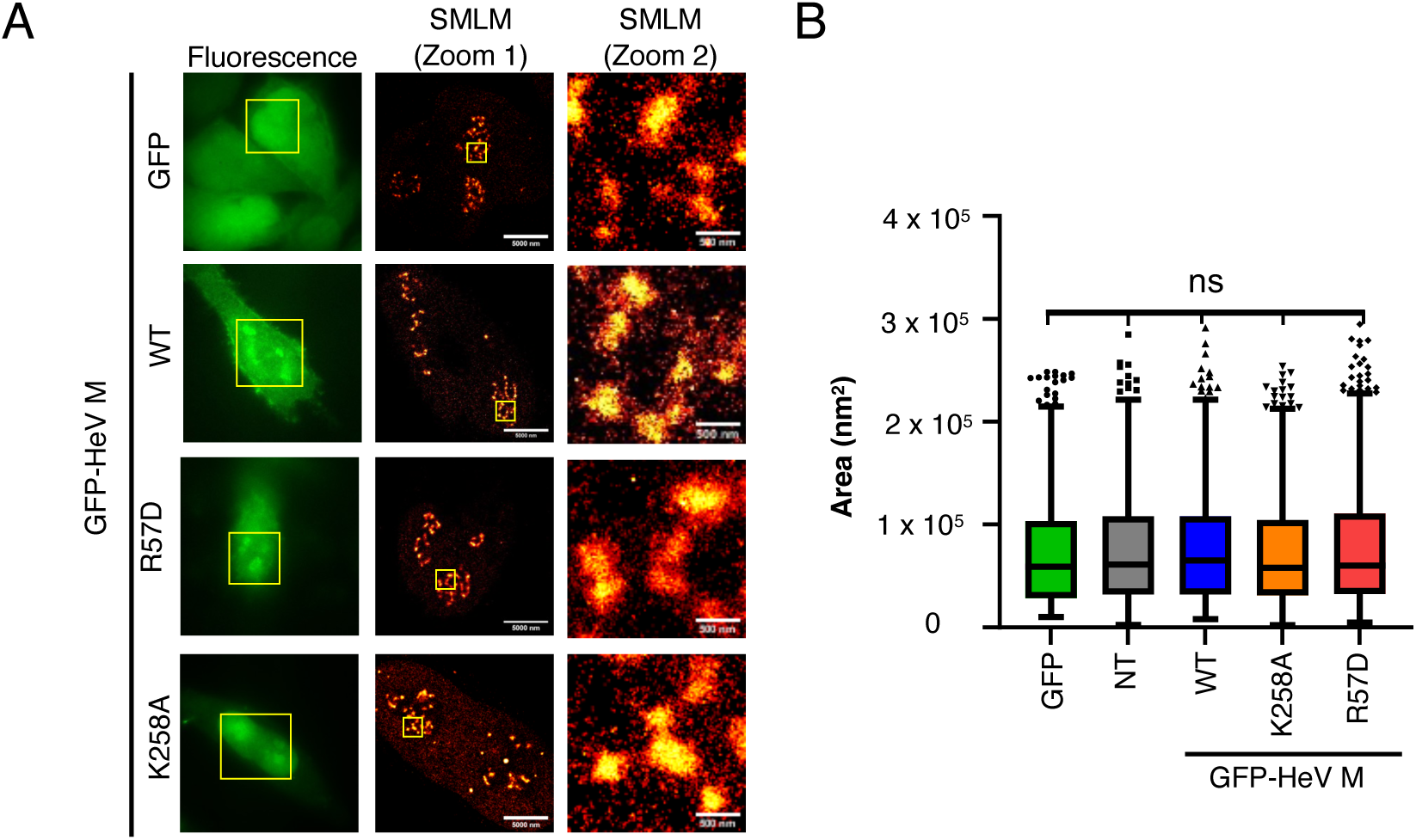
HeV M R57D protein does not affect the structure or distribution of Treacle-enriched FC-DFC. (A) HeLa cells expressing the indicated proteins or non-transfected (NT) were fixed and immunolabeled for Treacle before imaging using SMLM with *d*STORM. Fluorescence microscopy was used to identify cells expressing GFP/GFP-fusion proteins (green). Yellow boxes in fluorescence images show zoomed region imaged using SMLM (Zoom 1); yellow boxes in Zoom 1 images show additional zoomed region (Zoom 2). Scale bars correspond to 5000 nm (Zoom 1) and 500 nm (Zoom 2). SMLM images such as those shown in A were used to determine (B) Tukey boxplots showing median area and the 95% confidence interval of Treacle-enriched FC-DFC (n ≥ 505 FC-DFC for each sample). Statistical analysis used Kolmogorov-Smirnov tests as normality testing indicated a non-parametric distribution.

### Ubiquitination regulates sub-nucleolar trafficking of M proteins of multiple henipaviruses

Previously, we showed that the FC-DFC accumulation, Treacle binding, and inhibition of rRNA biogenesis are conserved among M proteins of multiple henipaviruses (including NiV, Cedar (CedV), and Mojiang (MojV) viruses), albeit with some differences in the extent of FC-DFC accumulation [23]. To determine if ubiquitin-dependence of FC-DFC accumulation is conserved in different henipaviruses, we assessed the effects of MG132 as above (e.g. Fig. 1). Similar to HeV M, MG132 treatment significantly impaired NiV M FC-DFC accumulation (Fig. 8A, B) and reduced the percentage of cells with FC-DFC accumulation (Fig. 8C) consistent with the homology of HeV and NiV M proteins (∼90% amino acid identity). Consistent with our previous report [23], CedV M showed the highest accumulation in FC-DFC and lowest GC accumulation of the M proteins assessed; F_FC-DFC/GC_ accumulation of CedV M protein was significantly reduced (but remained higher than that of HeV or NiV M proteins) following MG132 treatment, and the percentage of cells with FC-DFC accumulation of CedV M protein remained c. 100% (Fig. 8C). MojV M showed the lowest accumulation in FC-DFC (consistent with previous data [23]), resulting only a minor and non-significant reduction in F_FC-DFC/GC_ (Fig. 8B); however, there was a significant reduction in the percentage of cells with clear FC-DFC accumulation of MojV M protein following MG132 treatment (Fig. 8D).Taken together, these data indicate conserved roles of ubiquitination in regulating henipavirus M protein localization to the FC-DFC, although the extent of accumulation differs between M proteins, correlating with evolutionary divergence (c. 61% and 60% similarity of CedV and MojV M proteins, respectively, compared with HeV M protein).

**Figure 8.**
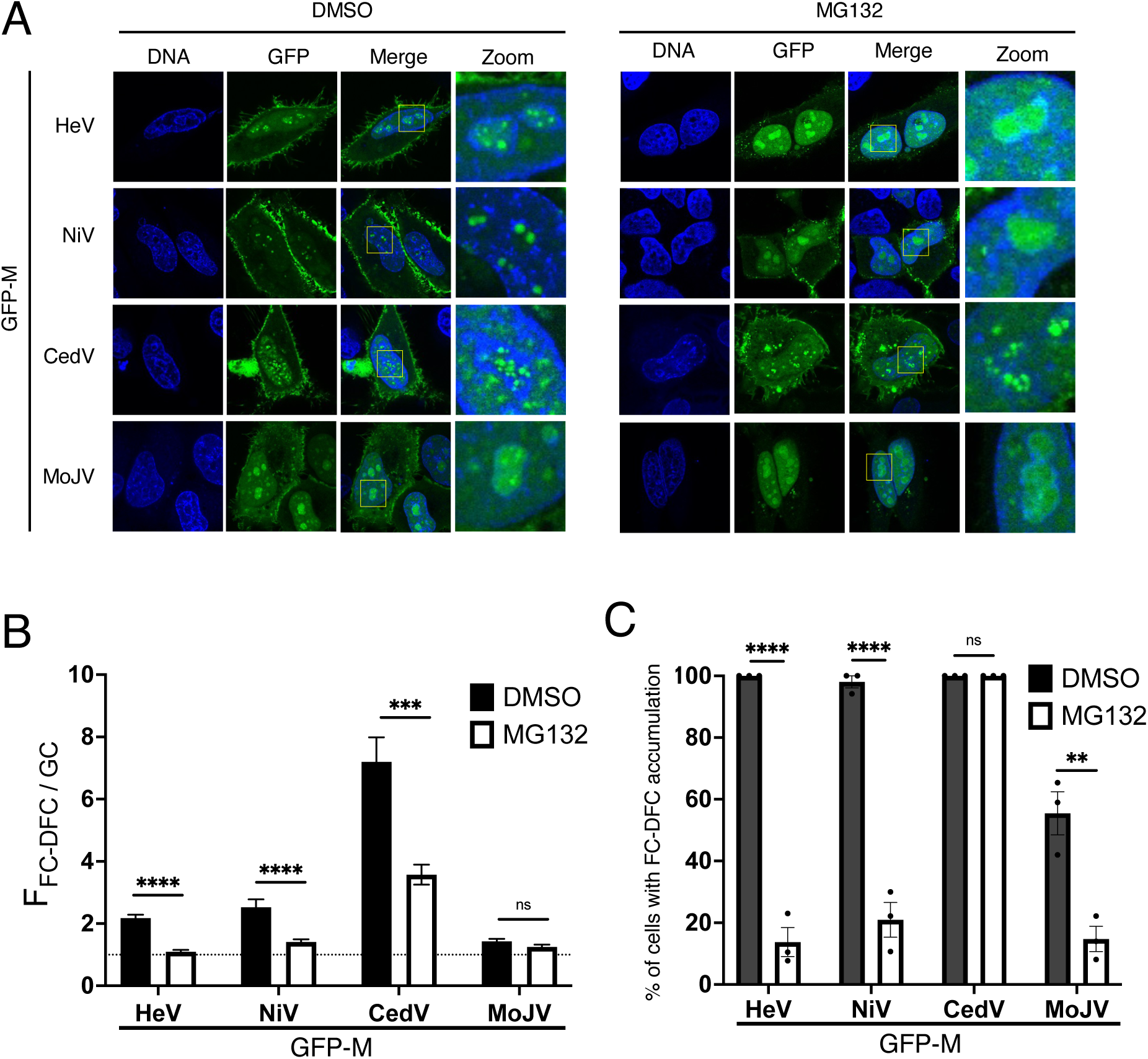
FC-DFC accumulation of M proteins of multiple henipaviruses is regulated by ubiquitination. (A) CLSM images of live HeLa cells transfected to express the indicated proteins. Representative images are shown for each condition, with yellow boxes magnified in the zoom panel. Hoechst 33342 was used to stain nuclei/DNA. (B) Images such as those in A were used to determine F_FC-DFC/GC_ (mean F_FC-DFC/GC_ ± S.E.M., from one assay (n ≥ 13), representative of three independent assays). (C) The percentage of M protein-expressing cells with apparent accumulation of M protein in FC-DFC (mean percentage ± S.E.M. from three independent assays; each sample was determined from n ≥ 13 cells).

## DISCUSSION

Here we have found that HeV M protein dynamically transits through the FC-DFC, indicating that the previously identified translocation through the nucleus/nucleolus involves additional sub-nucleolar trafficking between LLPS structures, with the different stages of trafficking regulated by post-translational modification. This transit enables regulated interactions with Treacle and other host factors, enabling functional regulation of rRNA synthesis by modulating the nucleolar DDR, as well as virus assembly and budding. To our knowledge, this study presents the first data on the mechanisms governing the trafficking of a viral protein within sub-nucleolar LLPS structures, and expands and refines the model for M protein trafficking. Specifically, we found that M trafficking to FC-DFC requires a basic residue at residue 258, and that its egress from FC-DFC requires oligomerization that is regulated, at least in part, by the ubiquitination status of the M protein. Importantly, while ubiquitination has previously been reported to regulate nuclear and overall nucleolar localization[10], indicating that ubiquitination is required for efficient nuclear export and nucleolar egress, our data, in contrast indicates that ubiquitination is required for nucleolar retention/accumulation into the FC-DFC. These findings highlight that localization of proteins to specific sub-nucleolar compartments involves highly specific mechanisms.

Taken together, our data support a model for M protein sub-nucleolar trafficking in which (1) entry of M protein dimers into the FC-DFC and functional interaction with Treacle to modulate rRNA biogenesis requires a basic charge at residue 258, and (2) exit from the FC-DFC requires oligomerization of the M protein within the FC-DFC. However, the retention/egress of M from the FC-DFC is dynamically regulated, at least in part, by the ubiquitination status of M protein. Our data support a likely role for ubiquitination of K258, but also indicate that other ubiquitination sites, either within M protein or on host proteins, also contribute, as FC-DFC localization by HeV M K258R protein is impaired but remains sensitive to inhibition of ubiquitination. These observations are consistent with previous data indicating that M protein can be mono-ubiquitinated at at least four sites, and K258R mutation inhibits ubiquitination at several of these sites (proposed to include K258 itself) which is likely to contribute to the impaired FC-DFC localization of this mutant. However, at least one site remains functional which may account for the residual accumulation of HeV M K258R into FC-DFC that is lost following MG132 treatment [10, 11]. Our data are consistent with a model whereby M protein exits the FC-DFC in the form of oligomers but egress is negatively regulated by ubiquitination of M protein; however, it is also possible that ubiquitination of host proteins also regulates interactions with M protein required for exit.

Movement of proteins between LLPS MLO structures such as nucleolar sub-compartments does not use conventional translocation processes associated with membrane-enclosed organelles (e.g. movement *via* pores/channels), but rather depends on partitioning through physicochemical properties and interactions with MLO-resident molecules [4, 26]. This likely accounts for the poor definition of nucleolar ‘targeting sequences’ compared with NLS and NES motifs that form specific interactions with trafficking receptor proteins (importins and exportins) [27]. Ubiquitination and oligomerization play important roles in the formation and regulation of LLPS [28, 29], and, notably, our data indicate a potential role for oligomerization, possibly regulated by ubiquitination, in coordinating M protein’s interactions/localization into sub-nucleolar liquid bodies. Thus, ubiquitination/oligomerisation may alter the physicochemical properties of M protein or interactions with constituents of different nucleolar condensates, as well as affecting importin/exportin interactions and interactions at budding sites. The differing nature of the mechanisms of sub-nucleolar trafficking and nucleocytoplasmic trafficking are consistent with our observations that ubiquitination has differing effects on exit from the FC-DFC to the GC, nucleolus to the nucleus, and nucleus to the cytoplasm. Thus, specific orchestration of ubiquitination, deubiquitination, oligomerization, and LLPS interactions may underlie appropriate temporal regulation of transport between these compartments, enabling specific control of rRNA silencing, virus replication, and assembly/budding, aligning with different stages of the viral life cycle [13].

As viruses typically mimic or hijack cellular processes, our findings likely have implications beyond viral infection. The intricate regulation of M protein in multiple intranuclear compartments is unlikely to have evolved solely to control the concentration in the cytoplasm for viral processes such as assembly and budding. Rather, it suggests a specific coordination of sub-nuclear functions, including DDR subversion, where M protein appears to mimic cellular NBS1 [14]. Our findings on the regulation of sub-nucleolar partitioning of M protein, including roles of positively charged residues (typical of nuclear/nucleolar targeting signals), ubiquitination and oligomerisation identifies mechanisms that may be relevant to cellular proteins that transit sub-nucleolar compartments, including those of the DDR. In the light of current advances toward therapeutic modulation of cellular and viral LLPS structures [30–33], our data also has the potential to contribute to novel antiviral approaches for currently incurable Henipavirus infection, and possibly other nucleolus-related pathologies [34, 35].

## MATERIALS AND METHODS

### Cell culture, transfection and treatment

HEK-293T (ATCC: CRL-3216) and HeLa (ATCC: CCL-2) cells were cultured in Dulbecco’s Modified Eagle Medium (DMEM) supplemented with 10% Fetal Calf Serum (FCS), 2 mM Glutamax, 50 U/mL Penicillin, and 50 μg/mL Streptomycin. The cells were maintained at 37 °C with 5% CO_2_. HEK-293T and HeLa cells were grown to 80-90% confluency before transfection using Lipofectamine 2000 and Lipofectamine 3000, respectively, according to the manufacturer’s instructions (ThermoFisher Scientific). For free ubiquitin depletion, transfected HeLa cells were treated with 50 μM MG132 or 0.5% DMSO for control at 18 h p.t. for 6 h before CLSM imaging analysis. MG132 was purchased from Sigma (M7449-200UL) as a 10 mM ready-made solution in DMSO. Experiments for co-expression of HA-Ubi, HeLa cells were transfected with the same total amount of DNA of 2500 ng: 1000 ng of GFP-HeV M co-transfected with 1500 ng total of HA-alone plasmid and/or HA-Ubi, with the HA/HA-Ubi ratio varied.

### Virus infections

Wild-type HeV (Hendra virus/horse/1994/Hendra) was used for all virus work and performed at the CSIRO Australian Centre for Disease Preparedness (CSIRO-ACDP) in Biosafety Level (BSL)-4 laboratories. For analysis of IF, HeLa cells were seeded onto coverslips and mock- or HeV-infected (MOI 5) prior to fixation at 7 h and 24 h p.i. using 4% paraformaldehyde (1h, RT) and permeabilization with 0.1% TritonX-100 for 10 min. IF labelling was performed using a mouse primary antibody to HeV M (1:500; developed internally (Ref#: 1805-21-1527) and an anti-mouse AlexaFluor 488 secondary antibody. DNA was visualised using a DAPI.

For tissue culture infective dose (TCID_50_) analysis HeLa cells were seeded into 96-well plates prior to HeV-infection the next day at MOI 0.5 or 5. At 18 h p.i. cells were treated with DMSO, MG132 (1 nM, 10 nM or 100 nM) or Bortezomib (5 nM, 50 nM or 500 nM). At 25 h p.i., additional Bortezomib was added to Bortezomib samples, as previously done for NiV[11]. At 42 h p.i. supernatants were collected and TCID_50_/ml was determined as previously[36].

### Constructs

Mammalian cell expression of N-terminal GFP tagged HeV-M (Accession Number AEB21196.1), and mutants were generated by directional cloning of the M gene cDNA into the multiple cloning site of the pEGFP-C1 vector, as previously described[17]. Site-directed mutagenesis of R57D residue was performed by applying QuickChange method, using *Pfu* DNA Polymerase from Promega. The following oligonucleotides were used to introduce the single R57D mutation: 5’-caagatctataccccaggtgcaaatgagGACaaattcaacaactacatgtacatg-3’ and 3’-catgtacatgtagttgttgaatttGTCctcatttgcacctggggtatagatcttg-5’. DNA sequencing was performed to confirm the correct mutation. The plasmid for expression of HA-ubiquitin (HA-Ubi) has been published previously [37].

### Confocal laser scanning microscopy (CLSM) and image analysis

For CLSM imaging analysis, HeLa cells were seeded on 1.5 (0.17 mm) thickness glass coverslips and transfected with the indicated constructs at 80-90% confluency. Imaging was performed at the indicated time p.t. or 24 h p.t., if not specified. CLSM was conducted using a Nikon C1 inverted confocal microscope with a 60× oil immersion objective (NA 1.4) at Monash Micro Imaging Facility. Live-cell CLSM imaging was performed within a heated chamber at 37 °C.

CLSM images were analyzed using ImageJ freeware software. The mean fluorescence of the nucleus (Fn), cytoplasm (Fc), nucleolus (Fnu; whole nucleolus), FC-DFC (F_FC-DFC_), GC (F_GC_), and background fluorescence (Fb) were determined. After subtracting the background fluorescence (Fb) from all values, the nuclear to cytoplasmic (Fn/c), nucleolar to nuclear (Fnu/n), and FC-DFC to GC (F_FC-DFC/GC_) fluorescence ratios were calculated. In cells where accumulation into sub-nucleolar compartments was not evident (e.g., cells expressing K258A- or K258R-mutated M protein), two distinct areas in the diffuse region of the nucleolus were selected to represent the “FC-DFC” and the “GC” for image analysis. The F_GC_ analysis was based on images captured under the same microscopy and software settings.

The percentage of cells with FC-DFC accumulation was determined by dividing the number of cells showing any nucleolus with FC-DFC ≥ 1 by the total number of cells expressing the indicated proteins in each sample. Data are presented as mean ± S.E.M. (standard error of the mean) or mean ± SD (standard deviation), as indicated in the figure legend. Statistical analysis (Student’s t-test) was performed using GraphPad Prism software.

### Immunofluorescence (IF)

For IF staining, cells grown on glass coverslips were washed twice gently with PBS at 24 h p.t., fixed with 4% (w/v) paraformaldehyde at room temperature (RT) for 15 min, permeabilized using 0.25% Triton X-100 (v/v in PBS) at RT for 5 min, and blocked with 1% bovine serum albumin (BSA) in PBS at RT for 1 h. For samples expressing FLAG-M proteins, an additional 5 min incubation with 5 μg/ml proteinase K was performed after fixation. The cells were then incubated with primary antibody specific to either Treacle (1:100; Cat # 11003-1-AP, Proteintech), UBF1 (1:500; Cat# Ab244287; Abcam), Nucleolin (1:200; Cat#14574, CST), anti-FLAG (1:250; Cat#F1804, Sigma) or NPM1 (1:200; Cat# 32-5200, ThermoFisher Scientific) at RT for 1.5 h. Subsequently, cells were incubated with goat anti-rabbit or anti-mouse 568 AlexaFluor conjugate secondary antibody (Cat # A-11011/A-11004, ThermoFisher Scientific) at a 1:1000 dilution in the dark at RT for 1.5 h. DNA staining was performed using Hoechst 33342 at a 1:2000 dilution of a 20 mM stock solution (Cat # 62249, ThermoFisher Scientific). The cells were mounted onto microscope glass slides (Lomb Scientific) using Mowiol reagent.

### 5-ethnyl uridine (EU) incorporation assays

Levels of rRNA synthesis were determined using an image-based technique (Click iT RNA Alexa Fluor 594 Imaging kit, Thermo-Fisher, Cat# C10330) as previously described [14, 23, 24, 38]. Cells were incubated for 1 h in the presence of EU before fixation in 4% paraformaldehyde at RT for 12 min and permeabilization in 0.25% Triton X-100 for 5 min at RT. Samples were processed according to the manufacturer’s recommendations to label incorporated EU with Alexa Fluor 594. DNA was labeled using Hoechst 33342. Cells were imaged by CLSM to detect labeling of nascent rRNA by measuring the fluorescence intensity of Alexa Fluor 594 within nucleoli. Quantitative analysis was performed using ImageJ software to determine the mean EU fluorescence of nucleoli, which were identified using a combination of DNA, GFP, and transmitted light channels. Relative rRNA synthesis levels were determined by measuring nucleolar EU levels of both GFP-expressing and non-expressing cells from the same sample (as an internal control) and expressed as nucleolar EU of GFP-expressing cells relative to non-expressing cells.

### Immunoprecipitation (IP) and immunoblotting (IB)

HEK-293T cells were seeded in 6 cm dishes and transfected to express the indicated GFP-fused proteins. At 24 h p.t., cells were harvested and lysed in 10 mM Tris/Cl pH 7.5, 150 mM NaCl, 0.5 mM EDTA, 0.5% NP-40, and 1x Protease Inhibitor Cocktail (PIC; Sigma-Aldrich Cat#11697498001) for 20 min at 4°C. The lysate was cleared by centrifugation at 20,000 xg for 6 min at 4°C, and 10% of the supernatant was collected as the ‘input’ sample. The remaining lysate was incubated with GFP-Trap magnetic beads (ChromoTek) as previously described for 45 min at 4°C on a rotary mixer [14]. The beads were then washed three times with dilution buffer (10 mM Tris/Cl pH 7.5, 150 mM NaCl, 0.5 mM EDTA, 1x PIC) and resuspended in 2x SDS-PAGE sample buffer to solubilize the proteins. The entire process from the addition of lysis buffer to the addition of 2x SDS-PAGE sample buffer was completed in less than 1.5 h to minimize degradation of Treacle, which is commonly observed (e.g. [24, 38, 39]).

Immunoprecipitates and cell lysates (input) were analyzed by SDS-PAGE and immunoblotting (IB). Proteins in IP and input samples were separated on 10% SDS-PAGE gels and transferred onto nitrocellulose membranes using the Bio-Rad Trans-Blot Turbo Transfer System according to the manufacturer’s instructions. After protein transfer, the membranes were blocked in blocking buffer (5% non-fat milk in PBS with 0.1% Tween-20) at RT for 1 h, incubated with the indicated primary antibodies at RT for 2 h or at 4°C overnight on a rocker, followed by incubation with HRP-conjugated secondary antibodies (goat anti-rabbit or anti-mouse) at RT for 1 h. The membranes were then imaged using a Gel Doc XR+ Gel Documentation System (Bio-Rad). The primary antibodies used for IB were anti-Treacle (Cat # 11003-1-AP, Proteintech; 1:2000), anti-GFP (Cat # 11814460001, Roche; 1:2000), and anti-β-tubulin (Cat # T8328, Sigma; 1:2000). The secondary antibodies used for IB were goat anti-rabbit (Cat # AP307P) and goat anti-mouse (Cat # AP308P) IgG HRP-conjugated antibodies (Merck Millipore). The Treacle bands in the IP were quantitated using ImageLab (Version 6.0) software.

### *d*STORM imaging and analysis

HeLa cells were fixed 24 h p.t. with 4% paraformaldehyde for 10 min then permeabilized with 0.1% Triton X-100 for 10 min. Fixed cells were blocked in 2% BSA/PBS for 30 min, then stained with anti-Treacle primary antibodies (3 µg/ml, 1 h) and Alexa Fluor 647 conjugated secondary antibodies (5 µg/ml, 45 min). Samples were imaged with a switching buffer of 100 mM mercaptoethylamine (MEA) in PBS made to pH 8.5 [40]. *d*STORM imaging was performed on a home-built super-resolution set-up as previously described [40, 41]. Briefly, the setup comprised an inverted fluorescence microscope (Olympus IX81, 100X 1.49 NA TIRF objective) with an EMCCD camera (Andor iXon). Blue laser excitation (Toptica iBeam 488 nm) was used to identify GFP-positive cells. High power red laser excitation (Oxxius LaserBoxx 638 nm) was used to induce reversible photoswitching of Alexa Fluor 647 fluorophores in the fixed samples. Single molecule ‘blinking’ events were captured at 100 Hz for between 10,000 – 20,000 frames and processed using rapi*d*STORM [42] to render a 2D coordinate map as the super-resolved *d*STORM images of subnucleolar Treacle. For analysis, images were first smoothed in ImageJ using the Gaussian Blur function (Sigma (Radius) = 0.5) to account for single molecule localization precision error. Images were converted to 8-bit (greyscale values 2-255 threshold) then converted into binary images. Using the “Distance Map” function in ImageJ, individual subnucleolar compartments were discretized based on pixel density by applying a threshold for greyscale values 2-255. The “Analyse Particles” function was then used to identify and measure subnucleolar compartments.

## ACKNOWLEDGEMENTS

We acknowledge the facilities and technical assistance of Monash Micro Imaging (Monash University) with confocal microscopy. This study was supported by National Health and Medical (NHMRC) Research Council grants 1125704, 1160838 and 1079211 to GWM; Australian Research Council (ARC) grants DP210100998, DP150102569 to GWM, and DP170104477 to TDMB; and the Miegunyah Trust Grimwade Fellowship to GWM.

