## Supplementary Figures for "Sub-nucleolar trafficking of Hendra virus matrix protein is regulated by ubiquitination and oligomerisation"

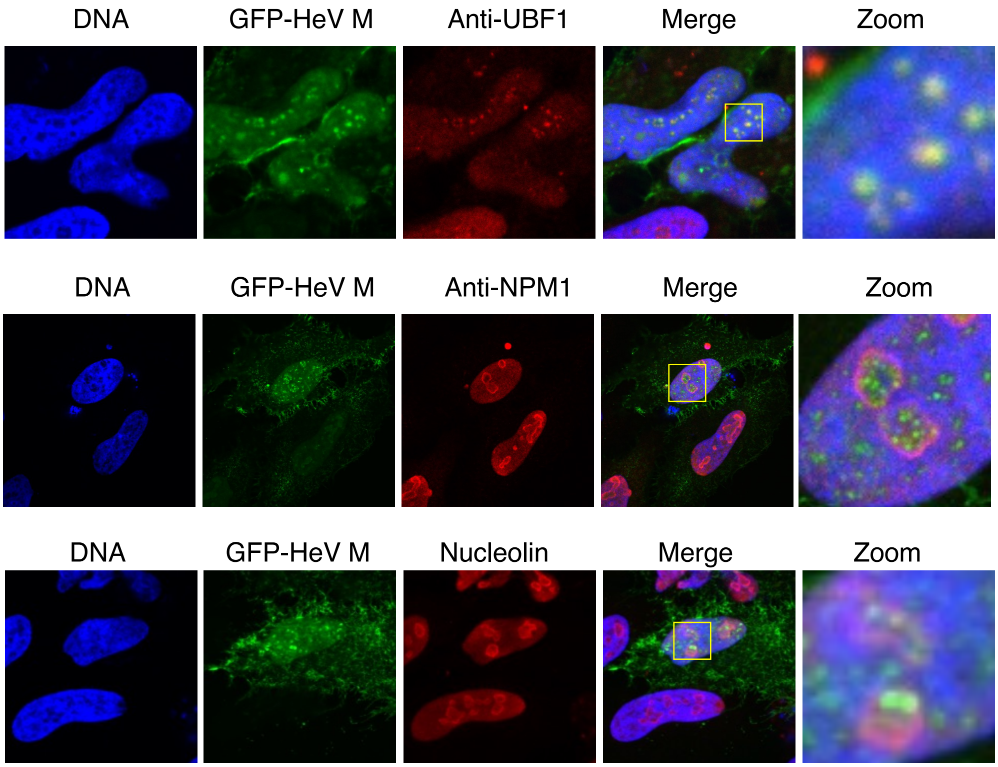


**Supplementary Figure 1. GFP-HeV M protein accumulation colocalizes with FC-DFC marker UBF1 but not GC markers.**

HeLa cells were transfected to express GFP-HeV M and fixed at 24 h p.t. with 4% paraformaldehyde before immunostaining for the nucleolar markers UBF1 (FC-DFC localization) and NPM1 and nucleolin (GC localization). Yellow boxes are magnified in the zoom panel. Hoechst 33342 (blue) was used to identify DNA/nuclei.

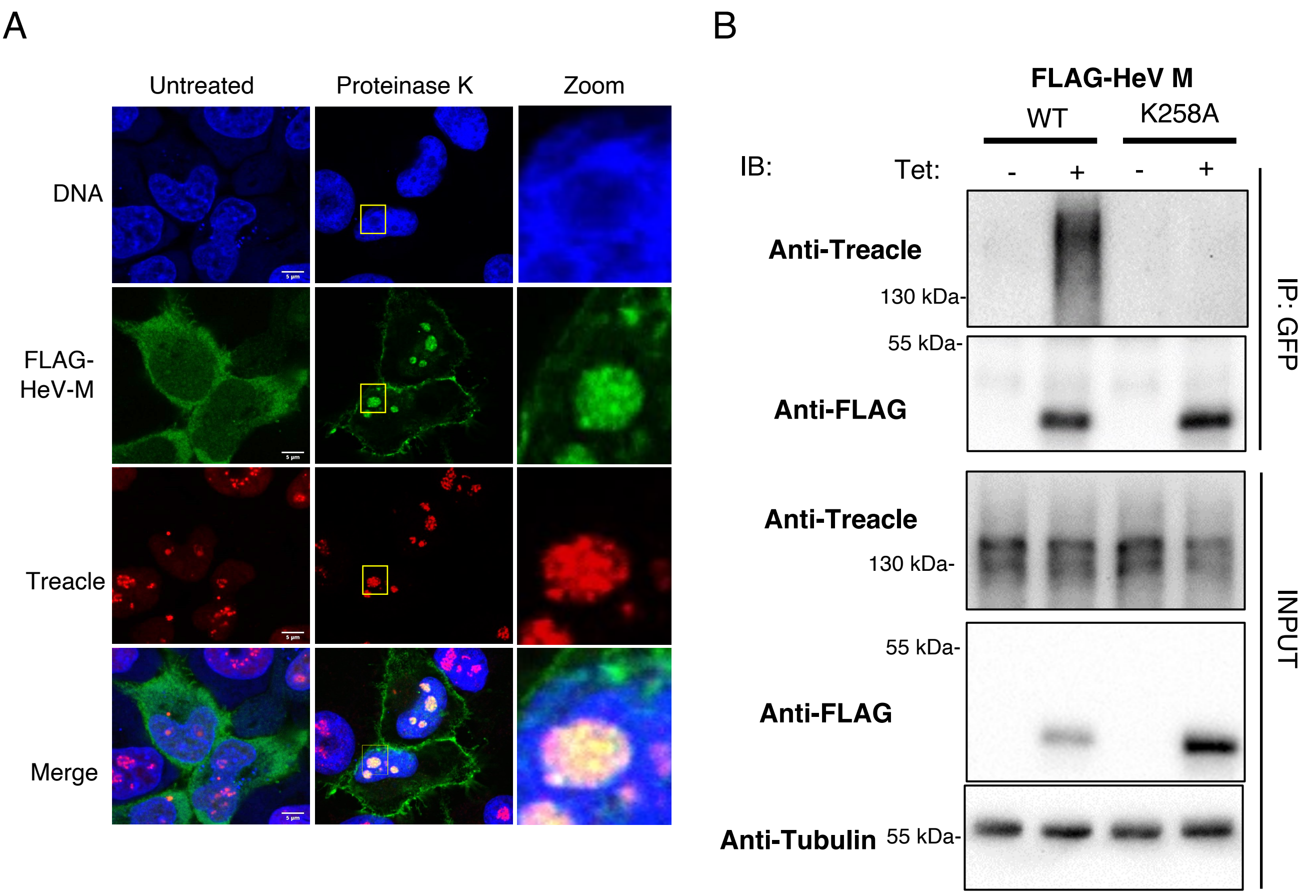


**<u>Supplementary Figure 2.</u> FLAG-HeV M binds Treacle and localizes to sub-nucleolar compartments.** (A) HeLa cells transfected to express FLAG-HeV M were fixed with 4% paraformaldehyde and treated with or without proteinase K, before immunostaining for FLAG (green) and Treacle (red). Yellow boxes are shown magnified in the zoom panel. Hoechst 33342 (blue) was used to identify DNA/nuclei. (B) 293 FLP-In™ cells stably transfected to enable inducible expression of 3xFLAG-HeV M or 3xFLAG-HeV M-K258A were treated with tetracycline (+Tet) to induce expressions, or not treated (-Tet), for 24 h before lysis and IP for FLAG; IPs were analysed by IB using the indicated antibodies.

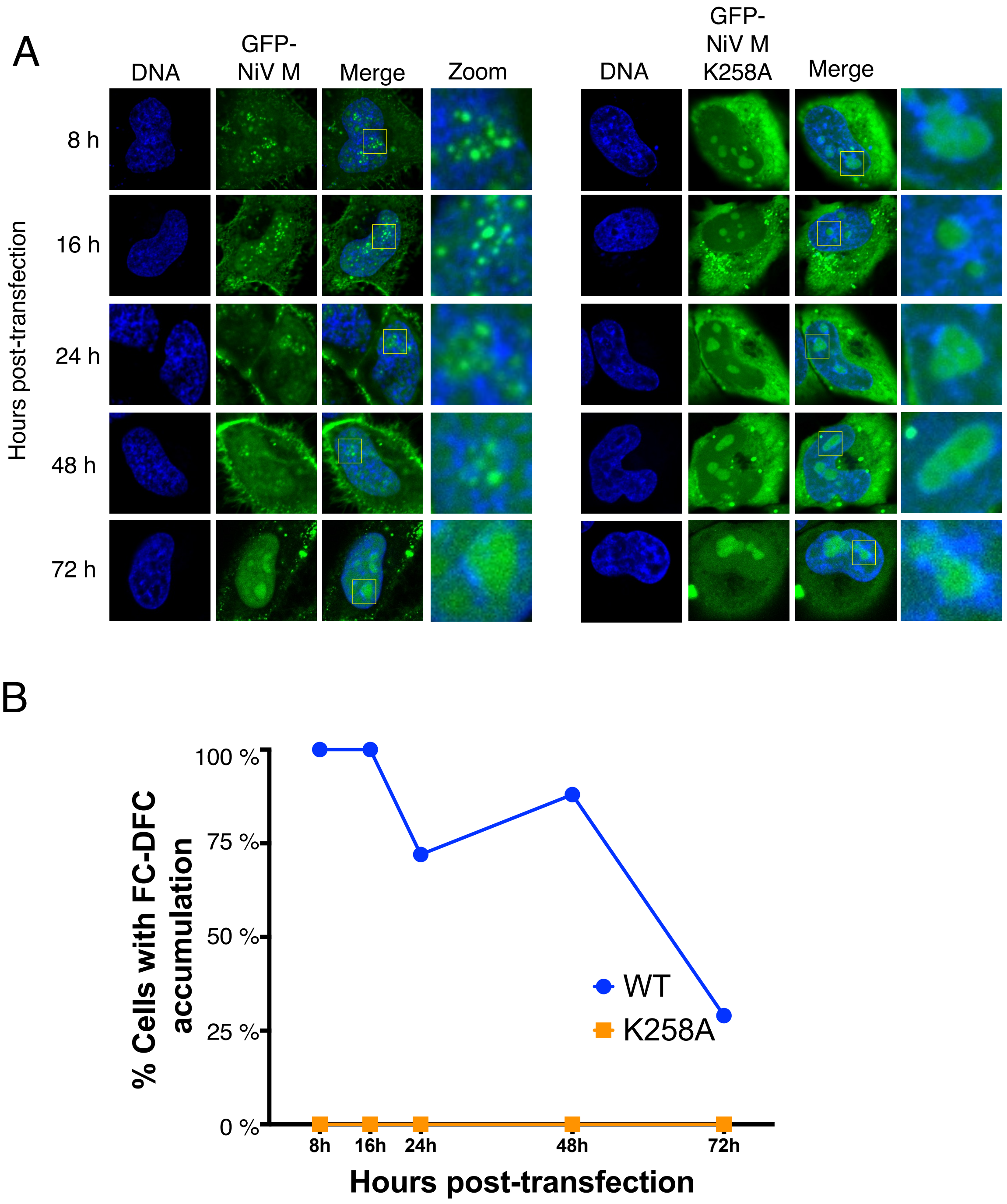


**Supplementary Figure 3.** **FC-DFC accumulation of NiV M protein decreases over time.**
(A) HeLa cells transfected to express GFP-NiV M WT or K258A proteins were analyzed live at 8, 16, 24, 48, and 72 h p.t. by CLSM. Images representative of major phenotypes are shown for each condition, with yellow boxes magnified in the zoom panel.
(B) Images such as those in (A) were used to determine the percentage of cells with apparent FC-DFC accumulation. The number of cells analyzed to determine the percentage for each sample is indicated on the graph. $n \geq 18$ cells were analysed for each time point.

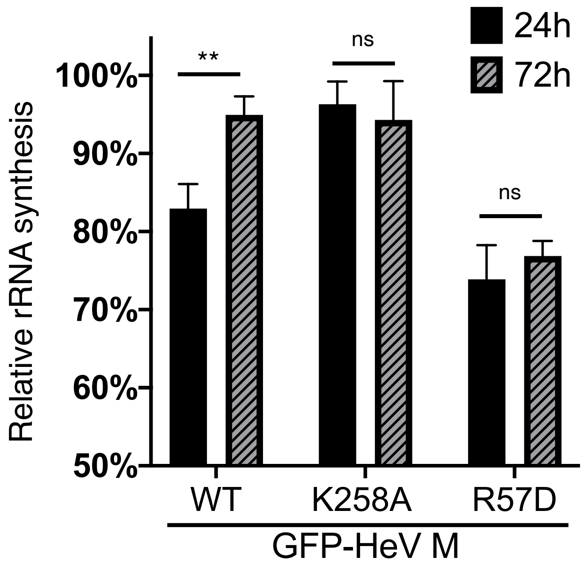


**Supplementary Figure 4. Inhibition of rRNA biogenesis by HeV M protein correlates with dynamic localization to FC-DFC.** HeLa cells were transfected to express the indicated proteins before addition of EU reagent. Sample were labeled to detect nascent RNA (EU fluorescence) and imaged by CLSM. EU fluorescence in the nucleoli of GFP-positive cells relative to that in non-GFP expressing cells in the same sample (mean relative EU fluorescence ± S.E.M., n $\geq 22$ cells) ** $p< 0.01$; ns, non-significant.
